# Implementation of a TMS-fMRI system: A primer

**DOI:** 10.1101/2021.05.19.444738

**Authors:** Golnoush Alamian, Ruiyang Ge, Afifa Humaira, Elizabeth Gregory, Erin L. MacMillan, Laura Barlow, Fidel Vila-Rodriguez

## Abstract

Transcranial magnetic stimulation (TMS) is a non-invasive and non-pharmacological intervention, approved for the treatment of individuals diagnosed with treatment-resistant depression. This well-tolerated approach uses magnetic pulses to stimulate specific brain regions and induce changes in brain networks at multiple levels of human functioning. Combining TMS with other neuroimaging techniques, such as functional magnetic resonance imaging (fMRI), offers new insights into brain functioning, and allows to map out the causal alterations brought on by TMS interventions on neural network connectivity and behaviour. However, the implemention of concurrent TMS-fMRI brings on a number of technical challenges that must be overcome to ensure good quality of functional images. The goal of this study was thus to investigate the impact of TMS pulses in an MR-environment on the quality of BRAINO phantom images, in terms of the signal of the images, the temporal fluctuation noise, the spatial noise and the signal to fluctuation noise ratio, at the University of British Columbia (UBC) Neuroimaging facility. The results of our analyses replicated those of previous sites, and showed that the present set-up for concurrent TMS-fMRI ensures minimal noise artefact on functional images obtained through this multimodal approach. This step was a key stepping stone for future clinical trials at UBC.

## 1. Introduction

### 1.1. Background

Psychiatric diseases such as major depressive disorder (MDD) are typically treated with a combination of psychotherapy and pharmaceutical treatments. This first response route however fails to improve the symptoms in about half of these patients. After two rounds of failed drug-trial treatment, affected individuals are categorized as having treatment resistant depression (TRD) (Conway et al., 2017). It is estimated that more than a third of MDD patients fail to achieve remission using antidepressants and are thus classified as having TRD (Nemeroff, 2007).

Over the years, this account has given way to alternative treatment options. Transcranial magnetic stimulation (TMS) (George et al., 1999; Hallett, 2000; Vila-Rodriguez et al., 2013), for instance, is a non-invasive and well-tolerated intervention, which has received both Health Canada and US FDA approval for the treatment of TRD. This approach relies on electromagnetic induction to stimulate brain regions focally, and induce changes in brain function at the cognitive, motor, emotional and behaviour levels. Specifically, TMS changes the brain’s electric current in target brain areas by sending magnetic pulses using a coil, which change the local field potentials at the surface of the cortex, thereby changing the excitatory/inhibitory neuronal activity within a given region (Downar et al., 2016). Of note, given its nature, TMS allows to investigate causality in the brain, by measuring functional changes associated with the electromagnetic stimulation, something that is inaccessible through other human brain mapping methods (Siebner et al., 2009). Hence, combining TMS interventions with a neuroimaging technique could reveal the critical temporal and spatial neural dynamics involved in the causal network changes that occur during TMS treatment in neurological and psychiatric patient populations.

One neuroimaging method that has long-been used for the study of brain activity and connectivity patterns is functional magnetic resonance imaging (fMRI). fMRI is a powerful tool that measures changes in the brain’s concentration of oxygenated and deoxygenated hemoglobin via the blood-oxygen-level-dependent (BOLD) signal (Huettel et al., 2004; Buxton, 2009). It is thought that the BOLD signal changes as a result of neuronal activity given that is coupled with cerebral blood flow and energy use (Logothetis, 2008). Moreover, given its’ precise millimetre spatial resolution and fair temporal resolution, fMRI is the most widely used neuroimaging technique to investigate brain function.

Concurrent TMS-fMRI is an emerging imaging approach that combines these two methodologies, and allows for TMS treatment inside the MR-scanner, while acquiring fMRI data. By employing this technique, cortical excitability can be manipulated, while monitoring the intra- and inter-cerebral functional activity of a brain region.

#### 1.1.1. Summary of previous TMS-fMRI experiments in healthy and clinical populations

Concurrent TMS-fMRI was first performed by (Bohning et al., 1997), and experiments have been on the rise in both healthy (Bohning et al., 1997; Nahas et al., 2001; Denslow et al., 2005; Bestmann et al., 2008b; de Vries et al., 2009; Moisa et al., 2009, 2010; Li et al., 2011; Ricci et al., 2012; Hanlon et al., 2013; Schintu et al., 2021; Oathes et al., 2021) and clinical populations (Li et al., 2004; Bestmann et al., 2006; de Vries et al., 2012; Webler et al., 2020). This is not surprising given that this multimodal approach allows to, not only examine the altered brain connectivity patterns of neurological and psychiatric patients but, also to investigate how local stimulation of focal brain regions can alter those same connectivity patterns, *in vivo*. This method can thus improve our overall understanding of brain functioning, while also refining where and how neurostimulation treatments, such as TMS, are offered to patient populations. Several reviews have assessed the capabilities of combining TMS with neuroimaging tools (Bohning et al., 2003a; Bestmann et al., 2008a; Siebner et al., 2009; Vila-Rodriguez et al., 2019; Bergmann et al., 2021), and discussed the implicated challenges and limitations. Among others, some of the advantages is the possibility to assess neuronal activity changes brought on by TMS using concurrent fMRI, mapping causal top-down changes and interactions, and investigating the causal relationship between brain region activations and behaviour (Bestmann et al., 2008a).

#### 1.1.2. What do we know about artefact control

Combining the physics of TMS and fMRI brings on a number of technical challenges that require consideration. One of these challenges is that the MRI scanner generates a stable and homogenous magnetic field, while the TMS apparatus uses an unstable and non-constant magnetic field. Therefore, in order to achieve proper concurrent TMS-fMRI, specialized equipment (e.g., MR-compatible TMS machine, and customized TMS and MR coils for concurrent TMS-fMRI) (Bohning et al., 2003b; Moisa et al., 2008; Navarro De Lara et al., 2015; Cobos Sánchez et al., 2020) and synchronization between MR sequences and TMS pulses (Bohning et al., 1999; Jung et al., 2016) is required. Some of the other technical challenges include static and dynamic artifacts. Static artefacts stem from the physical presence of the TMS apparatus in the MR scanner (Bungert and Bowtell, 2008; Bungert et al., 2012), while dynamic artifacts arise from using the TMS machine during and fMRI scan (e.g., radiofrequency, RF, noise; leakage currents) (Weiskopf et al., 2009).

### 1.2. Objective

Given the technical complexities involved in combining TMS with neuroimaging methods such as fMRI, it is important to investigate the dynamical artefacts that can be generated by using TMS in an MR environment, prior to leading concurrent TMS-fMRI experiments in human participants. Therefore, the objective of this study was to measure the effect of TMS pulses, concurrently with fMRI sequencing, on the quality of the fMRI images. Specifically, we examined the level of noise induced by the presence of TMS and by TMS pulses on the acquired images of a phantom at different levels of TMS pulse intensity. Although some studies have previously tackled this question at their own institution levels (Moisa et al., 2008; Andoh and Zatorre, 2012), replication at our own site, at the University of British Columbia (UBC), was necessary to ensure the validity and reliability of the quality of images produced by concurrent TMS-fMRI, prior to the start of clinical trials in patient populations.

## 2. Methods

### 2.1. Concurrent TMS-fMRI set up and stimulation protocol

This study was built on a standardized setup for concurrent TMS-fMRI at the UBC MRI centre. Echo-planar images were collected from a BRAINO phantom on a Philips Achieva 3.0 Tesla scanner. An array of two single channel SENSE Flex-L coils were used in place of a volume head coil to allow optimal positioning of TMS coil. To reduce any static artefact, a MR-compatible TMS coil was acquired (MRI-B91 non-ferromagnetic TMS figure-8 coil), along with a high-current filter box that blocked current leakage and filtered radio-frequency (RF) noise. In addition, as it has been suggested in the literature, the distance between the trigger and the MRI scanner was maximized by placing the TMS stimulator (MagPro X100) outside of the MR-room using an 8-meter extension cable. The set-up of the phantom and the TMS coil is shown in Figure 1B.

**Figure 1.**
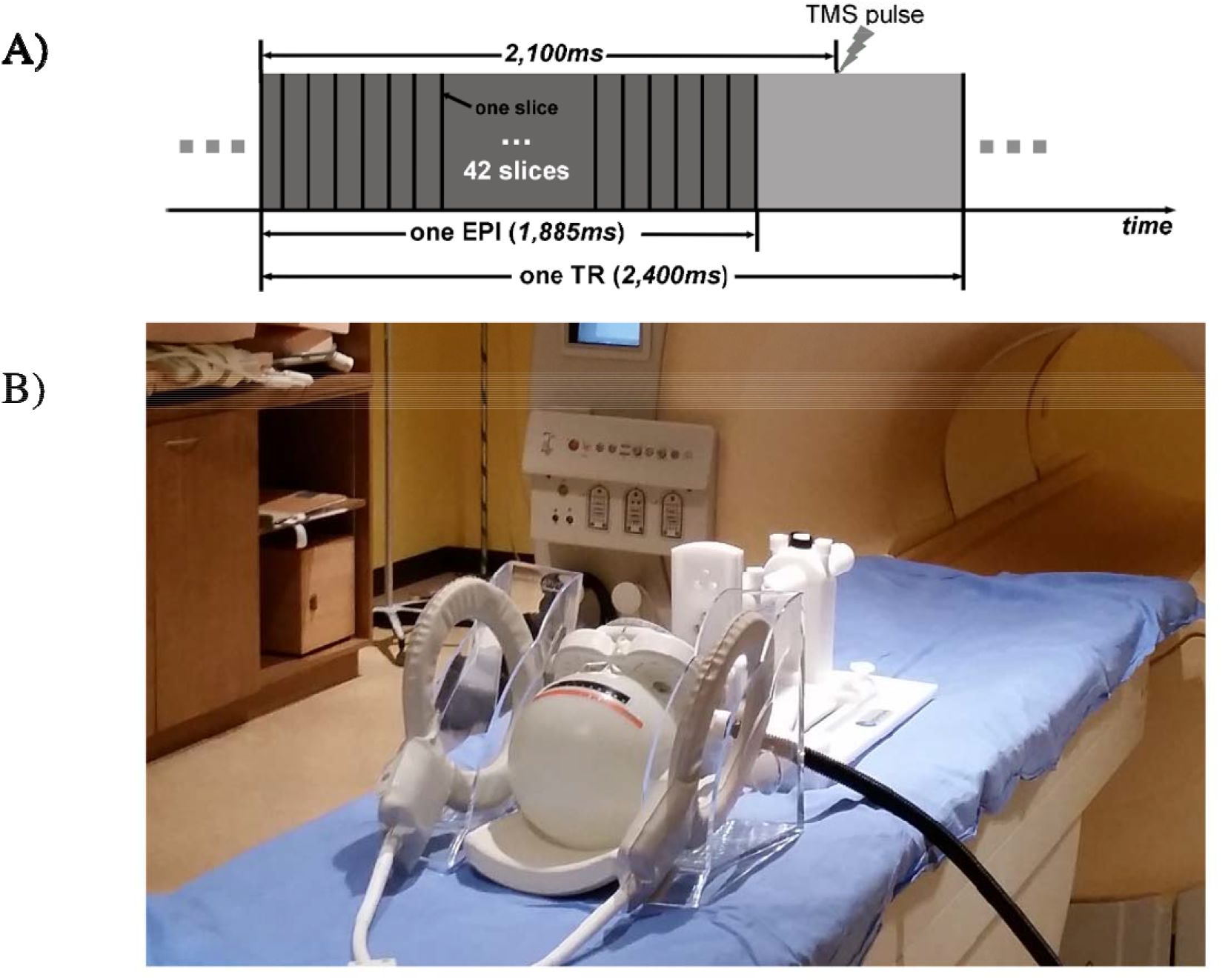
A) shows the stimulation protocol to test for potential artefacts when employing concurrent TMS and fMRI. B) illustrates the physical set-up of the phantom with the TMS coil inside the MRI scanner. EPI = echo planar imaging, fMRI = functional magnetic resonance imaging, TMS = transcranial magnetic stimulation, TR = repetition time

To determine the optimal TMS pulse firing time, an adjustable delay period was programmed into the synchronization hardware, which works as a relay between the TMS coil pulses and the MRI sequences. Specifically, the synchronization box first detects the single pulse generated by the MRI Philipps scanner at the beginning of each functional sequence. Next, the box amplifies this pulse to relay it to the TMS stimulator outside of the MRI room, in order to initiate a TMS pulse. During the concurrent TMS-fMRI scan, each echo-planar volume was acquired in 2400 ms. Previous literature has shown that if a period of at least 100 ms elapses after each TMS pulse, the subsequent MR images are distortion free (Bohning et al., 1999). Accordingly, the rTMS pulse was applied 2100 ms after the acquisition of the first slice, therefor allowing for an interval of 300 ms between the TMS pulse and the subsequent rs-MRI data acquisition (see Figure 1A). The synchronization between the TMS stimulator and echo planar image signal was achieved using an in-house developed software and synchronization hardware (partnership with PLC Electronics Solutions Ltd, Burnaby, Canada).

### 2.2. Stimulation protocol

The BRAINO spherical phantom was used to measure and compare the potential artefacts generated by the use of concurrent TMS-fMRI on the functional images. To evaluate image artefacts, the following conditions were examined: sequences (1) without TMS coil attached; (2) with TMS coil attached (in the zenithal position in direct contact with phantom), but with the stimulator turned off; (3) with TMS coil attached, and the stimulator turned to 0% of the maximum TMS stimulator output (i.e., 0% intensity); (4) with TMS coil attached, stimulating one pulse per time-repetition (i.e. one TMS pulse per MRI volume), and the stimulator turned to 30% intensity; and (5) with TMS coil attached, stimulating with one pulse per time-repetition, and the stimulator turned to 60% intensity. This sequential stimulation approach was taken to add one potential artefact factor with each condition, in order to better isolate and quantify the potential sources of image distortions. Specifically, condition (1) was the control condition, where any artefact generated at this stage was attributable to the MRI scanner noise itself. Condition (2) allowed to examine the potential artefact generated by the physical presence of the TMS coil in the MR-room. Condition (3) allowed to investigate the potential artefacts stemming from the TMS coil being energized. Finally, conditions (4) and (5) allowed to evaluate the effect of two different intensities of TMS pulses on the acquired images of the phantom.

### 2.3. Imaging Parameters and analyses

Each fMRI scans used a repetition time (TR) of 2400ms, a voxel size of 4mm × 4mm × 4.5mm, for a total of 210 phantom volumes, as shown in Figure 1A. To assess distortions on functional image qualities induced by using concurrent TMS-fMRI, several in-house Matlab scripts were developed.

Among the measures used to evaluate the quality of the images, first, the signal image, was computed by averaging the images obtained across the entire scanning session. Second, the temporal fluctuation noise image was calculated as the standard deviation of the residuals of the original, after subtracting the fit line (2^nd^ order polynomial). Third, the signal to fluctuation noise ratio (SFNR) was calculated by dividing the signal image by the temporal fluctuation noise image. Finally, the spatial noise image was derived via the difference of the averaged even-numbered fMRI volumes and the average of the odd-numbered fMRI volumes. SFNR summary values were computed as the average SFNR across voxels in the region of interest (ROI), for each condition, and was used to quantify the temporal noise of the fMRI data. In the present report, the ROIs included the entire phantom and its individual slices (see below). The analyses of the followed those of a previously validated report (Friedman and Glover, 2006).

## 3. Results

The results of the analyses of the image distortions due to the use of concurrent TMS-fMRI on a phantom are summarized in Figures 2 and 3. Figure 2 shows how the whole phantom image (the mean image of the 210 volumes acquired on each sequence), is altered across the five conditions in terms of the spatial noise and the temporal fluctuation noise. Overall, the images of the phantom did not show any obvious distortion under any of the five testing conditions, qualitatively. This held also true for the temporal fluctuation noise and the spatial noise.

**Figure 2.**
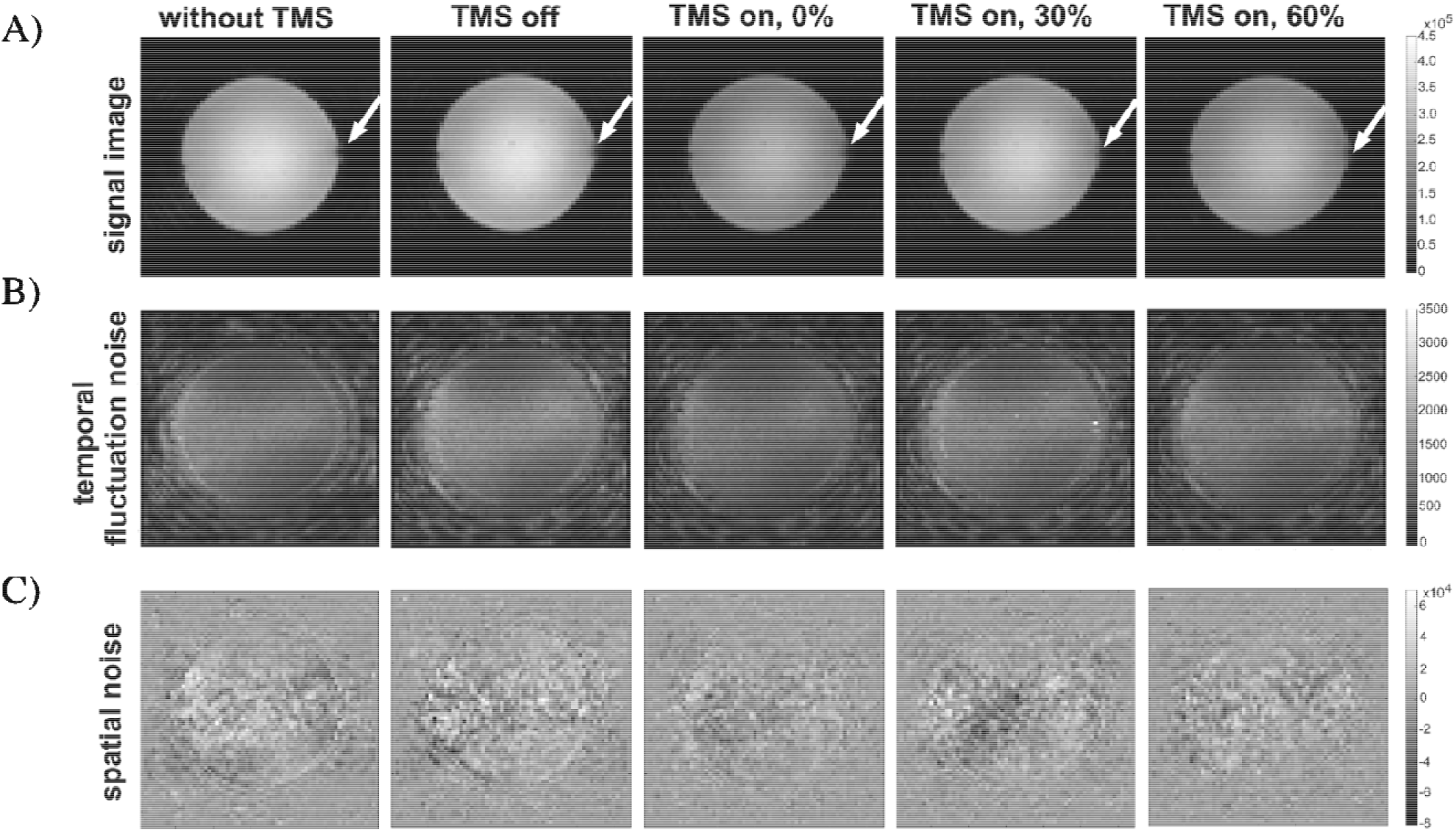
Signal and noise images for each test condition. A) shows the signal image and the arrow indicated the contact location between the phantom and the TMS coil. B) shows the spatial noise and C) shows the temporal fluctuation noise for each condition.

**Figure 3.**
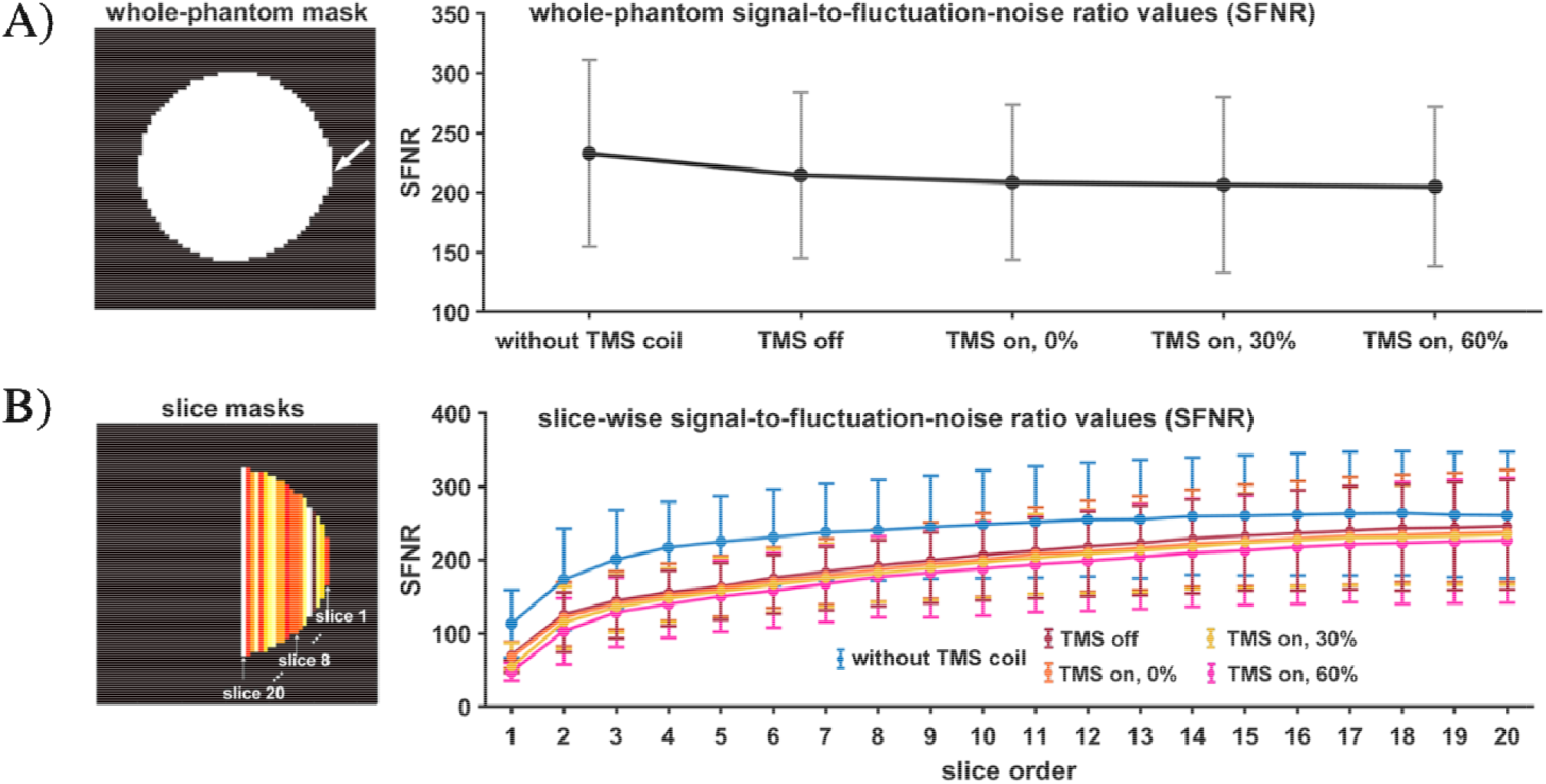
SFNR summary values (each slice as ROI) for each condition. A) The phantom mask in the left panel reveals the contact point between the phantom and the TMS coil (arrow) Moreover, we can see the change in SNFR as the condition changes, and as more potential artefact is added (from no TMS stimulation to TMS simulation at 60% intensity). Error bars indicate the standard deviation of the SFNR of the entire phantom voxels. B) shows how the SFNR changes in each condition for the first 20 slices of the masked phantom. Error bars indicate the standard deviation of the SFNR of all voxels included in the slice. SFNR=signal to fluctuation noise ratio, TMS = transcranial magnetic stimulation

Across all conditions, the whole-phantom SFNR summary value remained fairly stable, across the five testing conditions (Figure 3A). According to previous studies (Friedman and Glover, 2006), SFNR summary values of a regular fMRI image are typically around 150-200 in a 3T MRI scanner. In our experiment, the SFNR values across the five conditions ranged from 204.95 (condition 5) to 232.86 (condition 1), indicating that our fMRI data were of reasonable quality even with TMS firing on a relatively high intensity.

Next, we examined more closely how the SFNR changed in each condition, as a function to the proximity to the TMS coil. For Figure 3B, the ROI represents each 4mm slice, with slice 1 being the slice closest to the TMS coil, up to slice number 20, where the SFNR had stabilized. The results show that noise is maximal (low SFNR) at slice 1, near the TMS coil, and that it improves and stabilises by slice 4. To make a parallel with the human brain, slice 4 implies that we need to be approximately 16mm (slice 4, each slice 4mm thick = 16mm) away from the coil to obtain good SFNR. It has been estimated that the cortex lies approximately 10-20mm from the top of the scalp (Kozel et al., 2000; Beauchamp et al., 2011). Moreover, examining the changes per slices, Figure 3 B also illustrate that conditions 2-5 overlap significantly in terms of the slice-wise SFNR, and are slightly lower than those in the first condition, where the TMS apparatus is absent from the MR-room.

## 4. Discussion

This proof-of-concept phantom study demonstrated how concurrent TMS-fMRI procedures were implemented at the Neuroimaging facility of the University of British Columbia, for the purpose of using concurrent TMS-fMRI to better understand and treat TRD. Specifically, the use of an MRI-compatible TMS apparatus, a high-current filter box and physical separation of the TMS stimulator from the TMS coil helped reduce any potential static or dynamic artefacts that could have risen from the combination of these two neuroimaging tools (Ge et al., 2017). Given that the MRI scanner relies on a stable, constant, magnetic field, while the TMS apparatus relies on unstable magnetic field, we investigated where in the installation of the concurrent TMS-fMRI setup could potential artefacts arise from. To do so, a phantom was used to investigate when the TMS was not in the MRI room, when the TMS coil was in the MR-room but OFF, when the TMS coil was on ON, but at 0% pulse intensity, and when the TMS coil was turned ON to 30% and 60% intensity, respectively. The analyses of these sequential conditions revealed that minimal image distortion took place in conditions 2-5, implying that while noise was slightly increased due to the presence of the TMS coil and cable in the MR-room, the different intensities did not significantly affect the quality of the phantom volumes. Importantly, we observed that the SFNR increased and stabilized when the TMS coil was around 16mm from the site of interest, implying that temporal variability of the EPI signal reduced as the distance with the coil increased. As one would find it, this distance is approximately the distance between the human scalp and the cortex. This finding suggests that robust, low noise, signal can be obtained using concurrent TMS-fMRI at the cortical brain level. These findings replicate those other labs and researchers (Moisa et al., 2008; Andoh and Zatorre, 2012).

Despite these great strides, it is important to keep in mind that in human participants, TMS experiments can induce a startle or sensory response in the individual. These effects, and changes in the intensity or site of the stimulation during fMRI sequencing should be taken into consideration and investigated.

## 5. Conclusion

In sum, this study demonstrated the feasibility and artefact control involved in combining the spatial resolution of fMRI with the interventional functions of TMS at the University of British Columbia. This step was a critical step towards the clinical protocols aimed at treating TRD.

